# Poly(rC)-binding protein 1 limits hepatitis C virus virion assembly and secretion

**DOI:** 10.1101/2021.02.28.433252

**Authors:** Sophie E. Cousineau, Marylin Rheault, Selena M. Sagan

## Abstract

The hepatitis C virus (HCV) co-opts numerous cellular elements – including proteins, lipids, and microRNAs – to complete its viral life cycle. The cellular RNA-binding protein, poly(rC)-binding protein 1 (PCBP1), was previously reported to bind to the 5’ untranslated region (UTR) of the HCV genome; however, its importance in the viral life cycle has remained unclear. Herein, we sought to clarify the role of PCBP1 in the HCV life cycle. Using the HCV cell culture (HCVcc) system, we found that knockdown of endogenous PCBP1 resulted in an overall decrease in viral RNA accumulation, yet resulted in an increase in extracellular viral titers. To dissect PCBP1’s specific role in the HCV life cycle, we carried out assays for viral entry, translation, genome stability, RNA replication, as well as virion assembly and secretion. We found that PCBP1 knockdown did not directly affect viral entry, translation, RNA stability, or RNA replication, but resulted in an overall increase in infectious particle secretion. This increase in virion secretion was evident even when viral RNA synthesis was inhibited, and blocking virus secretion could partially restore the viral RNA accumulation decreased by PCBP1 knockdown. We therefore propose a model where endogenous PCBP1 normally limits virion assembly and secretion, which increases viral RNA accumulation in infected cells by preventing the departure of viral genomes packaged into virions. Overall, our findings improve our understanding of how cellular RNA-binding proteins influence viral genomic RNA utilization during the HCV life cycle.

## INTRODUCTION

Hepatitis C virus (HCV) is an enveloped virus of the *Flaviviridae* family (genus: hepacivirus) that typically causes a persistent liver infection (1). Its ~9.6 kb single-stranded, positive-sense RNA genome contains a single open reading frame flanked by 5’ and 3’ untranslated regions (UTR). A highly structured internal ribosomal entry site (IRES) in the 5’ UTR drives the translation of the viral polyprotein, which is subsequently processed into 10 mature viral proteins: 3 structural proteins (core, E1 and E2 glycoproteins), and 7 non-structural (NS) proteins (p7, NS2, NS3, NS4A, NS4B, NS5A and NS5B) (1, 2). While the structural proteins form the nucleocapsid and viral envelope, the NS3-5B proteins form the viral replicase, required for viral RNA replication (3). The p7, NS2, NS3 and NS5A proteins are also implicated in viral genome packaging and the assembly of viral particles (4–7). As a positive-sense RNA virus, the HCV genome itself must serve as a template for viral translation, genome replication, and packaging; however, the mechanisms that determine which process each viral RNA is engaged in at any given time have not been well defined. Importantly, due to their limited coding capacity, viruses are highly dependent on the molecular machinery of the host cell. While numerous cellular proteins and RNAs have been shown to interact with the HCV genome, their precise role(s) in the viral life cycle often remain poorly defined.

The poly(rC)-binding protein 1 (PCBP1), along with its paralogs hnRNP K and PCBP2, is one of the most abundant cellular RNA-binding proteins with an affinity for poly(rC) (8). These multifunctional proteins can regulate translation and enhance the stability of their cellular mRNA targets, which they interact with through their hnRNP K homologous (KH) domains (9). Notably, all three paralogs have been reported to bind to the HCV 5’ UTR (10–12). However, the degree to which each protein has been studied in the context of HCV infection varies significantly - while hnRNP K and PCBP2 have been fairly extensively studied and reported to play markedly different roles in the viral life cycle, the role of PCBP1 in the HCV life cycle has not been investigated in detail (13, 14). Beyond its interactions with the 5’ UTR, previous reports suggested that PCBP1 was not necessary for HCV IRES-mediated translation, but that knockdown of PCBP1 decreases HCV RNA accumulation during infection (15, 16).

Herein, we sought to further clarify the role of PCBP1 in the HCV life cycle. Using cell culture-adapted strains of HCV (HCVcc), we found that PCBP1 knockdown decreased viral RNA accumulation, yet led to an increase in virion secretion. By examining individual steps of the viral life cycle, we ruled out a direct role for PCBP1 in viral entry, translation, genome stability and viral RNA replication. Further analysis of the assembly step revealed that, similarly to its paralog hnRNP K, endogenous PCBP1 limits HCV virion assembly and release.

## MATERIALS AND METHODS

### Cell culture

Huh-7.5 human hepatoma cells were obtained from Charlie Rice (Rockefeller University) and maintained in complete media: Dulbecco’s Modified Eagle Media (DMEM) supplemented with inactivated 10% fetal bovine serum (FBS), 2 mM L-glutamine, and 1X MEM non-essential amino acids. Human embryonic kidney (293T) cells were kindly provided by Martin J. Richer (McGill University, Montreal, QC, Canada) and were maintained in DMEM supplemented with 10% FBS. All cell lines were maintained at 37°C/5% CO_2_ and were routinely screened for mycoplasma contamination.

### Plasmids and viral RNAs

The pJFH-1_T_ plasmid, encoding a cell culture-adapted Japanese Fulminant Hepatitis (JFH-1; HCV genotype 2a) with three adaptive mutations that increase viral titers in cell culture, was provided by Rodney Russell (Memorial University of Newfoundland) (17). The pJ6/JFH1 plasmid bears a full-length viral sequence derived from the J6 (structural genes through first transmembrane domain of NS2) and JFH-1 (remaining NS genes) isolates of HCV, and the pJ6/JFH-1 FL RLuc GNN (“RLuc-GNN”) viral sequence also includes a *Renilla* luciferase (RLuc) reporter gene inserted between p7 and NS2 and an inactivating GNN mutation within the NS5B RNA polymerase active site (4, 18). The pJ6/JFH-1 mono RLuc-NS2 plasmid (“Δcore-p7”) - a truncated version of the *Renilla* reporter virus with a deletion of the structural genes through p7 - was provided by Joyce Wilson (University of Saskatchewan) (19).

To make full-length uncapped viral RNAs, all plasmid templates were linearized and *in vitro* transcribed as previously described (20). The firefly luciferase (FLuc) mRNA was transcribed from the Luciferase T7 Control DNA plasmid (Promega) linearized using *XmnI* and *in vitro* transcribed using the mMessage mMachine T7 Kit (Life Technologies) according to the manufacturer’s instructions.

### Generation of infectious HCV stocks

To generate JFH-1_T_ stocks, 30 μg of *in vitro* transcribed RNA was transfected into Huh-7.5 cells using the DMRIE-C reagent (Life Technologies) according to the manufacturer’s instructions. Four days post-transfection, infectious cell supernatants were passed through a 0.45 μm filter and infectious viral titers were determined by focus-forming unit assay (17). Infectious virus was amplified for two passages through Huh-7.5 cells at a MOI of 0.2 before being used for infections. To generate J6/JFH-1 stocks, 25 μg of *in vitro* transcribed RNA was electroporated into 6 x 10^6^ Huh-7.5 cells using a Bio-Rad GenePulser XCell at 270 V, 950 μF, and infinite resistance in a 4-mm cuvette. Two days post-electroporation, infectious cell supernatants were passed through a 0.45 μm filter and concentrated at least 10-fold using Amicon Ultra-15-100K centrifugal filter units (Millipore Sigma) according to the manufacturer’s instructions. Viral titers were determined by focus-forming unit assay. All viral stocks were aliquoted and stored at −80°C until use.

### Focus-forming unit (FFU) assays

One day prior to infection, 8-well chamber slides (Lab-Tek) were seeded with 4 x 10^5^ Huh-7.5 cells/well. Infections were performed with 10-fold serial dilutions of viral samples in 100 μL for 4 h, after which the supernatant was replaced with fresh media. Three days post-infection, slides were fixed in 100% acetone and stained with anti-HCV core antibody (1:100, clone B2, Anogen), and subsequently with the AlexaFluor-488-conjugated anti-mouse antibody (1:200, ThermoFisher Scientific) for immunofluorescence analysis. Viral titers are expressed as the number of focus-forming units (FFU) per mL.

Extracellular virus titers were determined directly from cell supernatants, while intracellular virus titers were determined after cell pellets were subjected to lysis via four freeze-thaw cycles, removal of cellular debris via centrifugation, and recovery of virus-containing supernatants.

### MicroRNAs and siRNA sequences

siGL3 (siCTRL): 5’-CUUACGCUGAGUACUUCGAUU-3’, siGL3*: 5’-UCGAAGUACUCAGCGUAAGUU-3’, miR122_p2-8_ (siCTRL for luciferase experiments): 5’-UAAUCACAGACAAUGGUGUUUGU-3’, miR122_p2-8_*: 5’-AAACGCCAUUAUCUGUGAGGAUA-3’ (21), siPCBP1: 5’-CUGUGUAAUUUCUGGUCAGUU-3’, siPCBP1*: 5’-CUGACCAGAAAUUACACAGUU-3’ (16) were all synthesized by Integrated DNA Technologies.

All microRNA and siRNA duplexes were diluted to a final concentration of 20 μM in RNA annealing buffer (150 mM HEPES pH 7.4, 500 mM potassium acetate, 10 mM magnesium acetate), annealed at 37°C for 1 h and stored at −20°C. For all knockdown experiments, 50 nM siRNA transfections were conducted 2 days prior to infection or electroporation of viral RNAs. Transfections were conducted using Lipofectamine RNAiMAX (Invitrogen) according to the manufacturer’s instructions with the modification that 20 μL of reagent were used to transfect a 10-cm dish of cells.

### HCV and VSV pseudoparticles (HCVpp and VSVpp)

HCVpp consisting of a FLuc reporter lentiviral vector pseudotyped with the HCV E1 and E2 glycoprotein (from the H77 isolate, a genotype 1a strain) were a kind gift from John Law (University of Alberta) (22). To generate lentiviral vectors pseudotyped with the VSV-G glycoprotein (VSVpp), a 90% confluent 10-cm dish of 293T cells were transfected with 10 μg pPRIME-FLuc, 5 μg psPAX.2, and 2.5 μg pVSV-G plasmid with 10 μL Lipofectamine 2000 (Invitrogen) diluted in 4 mL Opti-MEM. Media was changed 4, 20, and 28 h post-transfection. At 48 h post-transfection, the cell culture media was passed through a 0.45 μm filter and stored at −80°C.

To assay for cell entry, HCVpp and VSVpp were diluted 1/3 in dilution media (1X DMEM, 3% FBS, 100 IU penicillin and 100 μg/mL streptomycin) with 20 mM HEPES and 4 μg/μL polybrene, and introduced to Huh-7.5 cells by spinoculation at 1,200 rpm for 1 h at room temperature. The cells were left to recover at 37°C for at least 5 h before the pseudoparticle-containing media was changed for fresh complete Huh-7.5 media. In parallel, cells seeded in a 6-well plate were transfected with 1 μg of pPRIME-FLuc plasmid using Lipofectamine 2000 (Invitrogen) according to the manufacturer’s instructions. Three days post-spinoculation and transfection, cells were lysed in passive lysis buffer (Promega) and FLuc activity was assayed using the Dual Reporter Luciferase kit (Promega).

### Infections

Three days prior to infection, 10-cm dishes were seeded with 5 x 10^5^ Huh-7.5 cells, which were transfected with siRNA duplexes on the following day. On the day of infection, each 10-cm dish - containing approximately 1 x 10^6^ cells - was infected with 5 x 10^4^ FFU of JFH-1_T_ diluted in 3 mL complete media. Four to five hours post-infection, each infected plate was split into three 10-cm dishes. Protein, RNA, and virus samples were collected three days post-infection.

### Electroporations

For each electroporation, 400 μL of resuspended cells (1.5 x 10^7^ cell/mL) were mixed with 2 μg of FLuc mRNA and 5 μg of replicating (Δcore-p7 J6/JFH-1 RNA) or 10 μg GNN J6/JFH-1 RNA, and electroporated in 4-mm cuvettes at 270 V, 950 μF, and infinite resistance, optimized for the Bio-Rad GenePulser XCell (Bio-Rad). Electroporated cells were resuspended in complete Huh-7.5 media and transferred to 6-well plates for luciferase assays and protein analysis.

### Inhibition of RNA replication by 2’CMA

Two days post-siRNA transfection, Huh-7.5 cells were infected with JFH-1_T_ at a MOI of 0.05. Four to five hours post-infection, each plate of infected cells was split into 6-well plates. Three days post-infection, the media on these cells was changed for complete Huh-7.5 media with 5 μM 2’CMA (2’C-methyladenosine, Carbosynth), an HCV NS5B polymerase inhibitor, or DMSO (vehicle control) (23). Total RNA and intracellular virus samples were collected at 0, 6 and 12 h post-treatment, while cell culture supernatants were collected 6 and 12 h post-treatment. Protein samples were collected from untreated plates to assess PCBP1 knockdown efficiency by Western blot.

### Inhibition of virus secretion by Brefeldin A

Three days post-infection with JFH-1_T_, the cell culture media was replaced with complete Huh-7.5 media with 5 μg/mL BFA (Brefeldin A, Biolegend) or equal volumes of DMSO (vehicle control) (24). Total RNA, intracellular virus, and extracellular virus samples were collected at 0 and 6 h post-treatment. Protein samples were collected from untreated cells to verify PCBP1 knockdown by Western blot.

### Western blot analysis

To collect total intracellular protein samples, cells were lysed in RIPA buffer (150 mM sodium chloride, 1% NP-40, 0.5% sodium deoxycholate, 0.1% SDS, 50 mM Tris pH 8.0), supplemented with Complete Protease Inhibitor Cocktail (Roche) and frozen at −80°C. Cellular debris was pelleted by centrifugation at 16,000 x g for 30 min at 4°C, and the supernatant was quantified by BCA Protein Assay (ThermoScientific). Ten micrograms of sample were loaded onto 10-12% SDS-PAGE gels. Samples were transferred onto Immobilon-P PVDF membranes (Millipore), blocked in 5% milk, and incubated overnight with primary antibodies diluted in 5% BSA: rabbit anti-PCBP1 (clone EPR11055, Abcam ab168378, diluted 1:10,000); rabbit anti-actin (A2066, Sigma, 1:20,000); mouse anti-HCV core (clone B2, Anogen MO-I40015B, 1:7,500); mouse anti-JFH-1 NS5A (clone 7B5, BioFront Technologies, 1:10,000). Blots were incubated for 1 hour with HRP-conjugated secondary antibodies diluted in 5% skim milk: anti-mouse (HAF007, R&D Systems, 1:25,000); anti-rabbit (111-035-144, Jackson ImmunoResearch Laboratories, 1:50,000) and visualized using enhanced chemiluminescence (ECL Prime Western Blotting Detection Reagent, Fisher Scientific).

### RNA isolation and Northern blot analysis

Total RNA was harvested using TRIzol Reagent (ThermoFisher Scientific) according to the manufacturer’s instructions. Ten micrograms of total RNA were separated on a 1% agarose gel containing 1X 3-(N-morpholino)propanesulfonic acid (MOPS) buffer and 2.2 M formaldehyde and transferred to a Zeta-probe membrane (Bio-Rad) by capillary transfer in 20X SSC buffer (3 M NaCl, 0.3 M sodium citrate). Membranes were hybridized in ExpressHyb Hybridization Buffer (ClonTech) to random-primed ^32^P-labeled DNA probes (RadPrime DNA labelling system, Life Technologies) complementary to HCV (nt 84-374) and γ-actin (nt 685-1171). Autoradiograph band densities were quantified using Fiji (25).

### RT-qPCR analysis

The iTaq Universal Probes One-Step kit (Bio-Rad) was used to perform duplex assays probing for the HCV genome (NS5B-FW primer: 5’-AGACACTCCCCTATCAATTCATGGC-3’; NS5B-RV primer: 5’-GCGTCAAGCCCGTGTAACC-3’; NS5B-FAM probe: 5’-ATGGGTTCGCATGGTCCTAATGACACAC-3’) and the GAPDH loading control (PrimePCR Probe assay with HEX probe, Bio-Rad). Each 20 μL reaction contained 500 ng of total RNA, 1.5 μL of the HCV primers and probe, and 0.5 μL of the GAPDH primers and probe. RT-PCR reactions were conducted in a CFX96 Touch Deep Well Real-Time PCR system (Bio-Rad). Genome copies were calculated using a standard curve and fold-differences in gene expression were calculated using the 2^-ΔΔCt^ method (26).

### Luciferase assays

For translation and replication assays, cells were washed in PBS and harvested in 100 μL of 1X passive lysis buffer (Promega). The Dual-Luciferase Assay Reporter Kit (Promega) was used to measure both *Renilla* and firefly luciferase activity according to the manufacturer’s instructions with the modification that 25 μL of reagent were used with 10 μL of sample. All samples were measured in triplicate.

### Data analysis

Statistical analyses were performed using GraphPad Prism v9 (GraphPad, USA). Statistical significance was determined by paired t-test to compare results obtained from multiple experiments, and by two-way ANOVA with Geisser-Greenhouse and Bonferroni corrections when more than two comparisons were applied at once. To calculate half-lives, a one-step decay curve using the least-squares regression was used, and error was reported as the asymmetrical (profile likelihood) 95% confidence interval of the half-life. To calculate virus accumulation and virus secretion rates, a simple linear regression was performed using the least squares regression method. The slope and standard error calculated for each regression represents the rate of virus accumulation or secretion.

## RESULTS

### PCBP1 plays a role in the HCV life cycle

Previous reports suggested that the PCBP1 protein directly interacted with the 5’ UTR of the HCV genome, and a siRNA screen reported that PCBP1 knockdown reduced HCV RNA accumulation (10, 16). Thus, we sought to further characterize the role of PCBP1 in the HCV life cycle. Firstly, we assessed how PCBP1 knockdown affected the accumulation of HCVcc using two viral strains: the cell culture-adapted JFH-1_T_ strain (which has three adaptive mutations in E2, p7, and NS2), and the chimeric J6/JFH-1 strain (where the coding region of the core through to the first transmembrane domain of NS2 comes from the J6 sequence, and the remaining sequences are derived from JFH-1) (**Figure 1A**) (17, 18). We found that knocking down endogenous PCBP1 decreased viral protein expression and HCV RNA accumulation of both strains, resulting in an approximately 2.2-fold and 3.3-fold decrease in JFH-1_T_ and J6/JFH-1 viral RNA accumulation, respectively (**Figure 1B-D**). Quantification of intracellular virions revealed no significant differences in JFH-1_T_ titers between the PCBP1 knockdown and control conditions; however, we observed an approximately 2.6-fold reduction in intracellular J6/JFH-1 titers during PCBP1 knockdown (**Figure 1E**). Surprisingly, when we quantified extracellular virions, we observed that PCBP1 knockdown resulted in an overall increase in extracellular titers of approximately 3.9-fold and 2.6-fold for JFH-1_T_ and J6/JFH-1, respectively (**Figure 1F**). Overall, PCBP1 knockdown increased the total quantity of virions produced by 3.7-fold for JFH-1_T_ and 2.1-fold for J6/JFH-1 (**Supplementary Figure S1**). To verify that this effect was specifically due to the decrease in endogenous PCBP1 protein expression, we attempted to rescue PCBP1 protein levels by ectopically expressing a siRNA-resistant PCBP1 with a N-terminal FLAG tag (**Supplementary Figure S2**). Although we were unable to ectopically express PCBP1 to the same level as the endogenous protein, we observed that ectopic PCBP1 expression diminished the impact of endogenous PCBP1 knockdown. Thus, in line with previous findings, we found that PCBP1 knockdown decreased viral protein expression and intracellular viral RNA accumulation. However, despite this overall decrease in viral protein and RNA accumulation, we observed a marked increase in virion production and these effects were partially reversed by ectopic expression of PCBP1.

**Figure 1.**
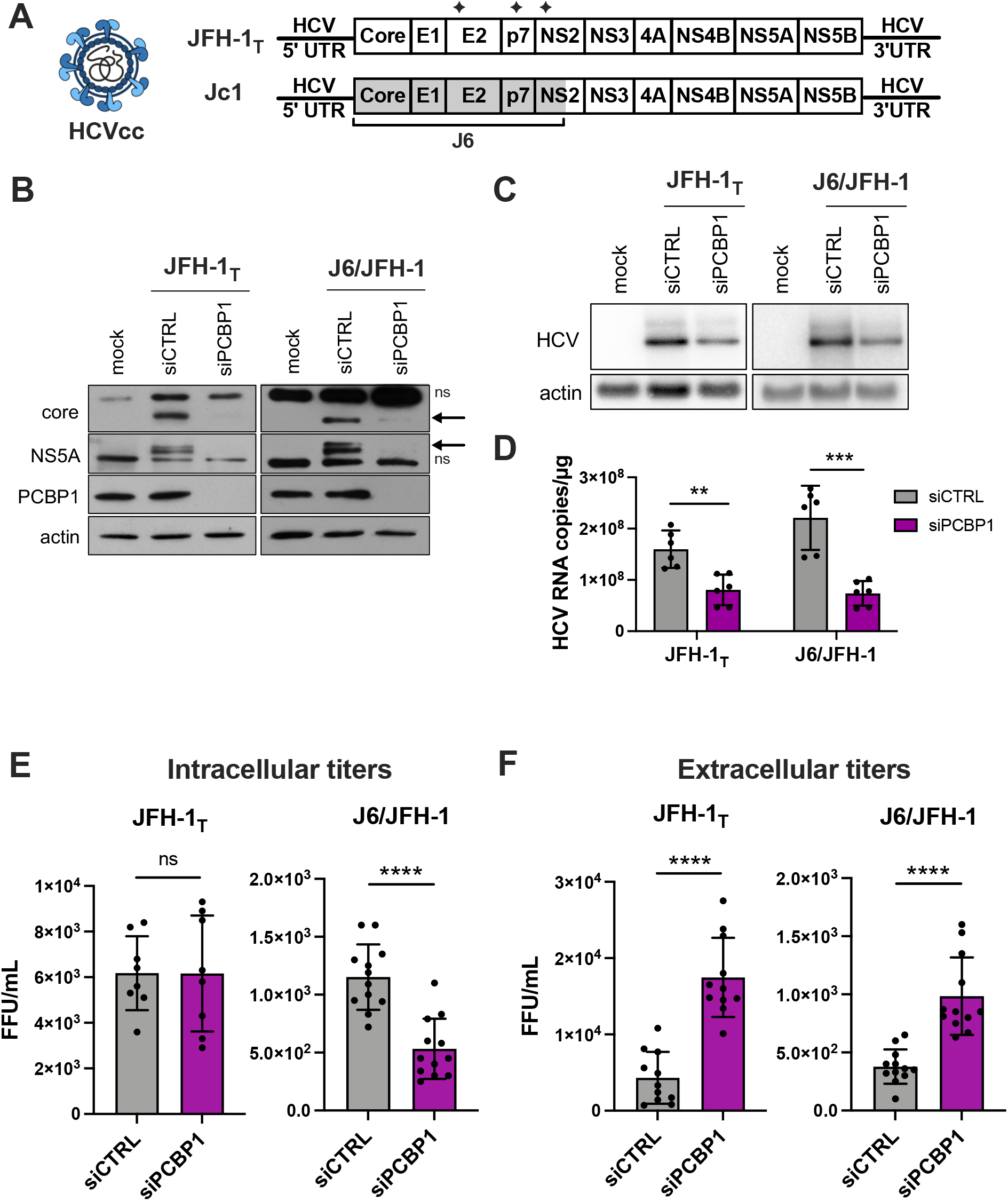
PCBP1 knockdown decreases HCV protein expression and intracellular viral RNA accumulation, but increases secreted virus titers. **(A)** Schematic representation of the HCVcc infectious particles and genomic RNA used in infections. Huh-7.5 cells were transfected with siPCBP1 or siCTRL two days prior to infection with JFH-1_T_ or J6/JFH-1 (MOI = 0.05). Total protein, RNA, and intracellular and extracellular infectious virus were harvested at day 3 post-infection. **(B)** Viral protein expression analysis by Western blot (ns, non-specific band; arrow, core and NS5A bands). **(C)** Viral RNA accumulation analysis by Northern blot and **(D)** quantification by RT-qPCR. **(E)** Intracellular and **(F)** extracellular (secreted) virus titers, quantified by FFU assay. All data are representative of at least three independent biological replicates, and error bars represent the standard deviation of the mean. Statistical significance was calculated by paired t-test (ns, not significant; ** p < 0.01; *** p < 0.001; **** p < 0.0001).

### PCBP1 knockdown does not have a direct impact on HCV entry, translation, genome stability or viral RNA replication

To clarify PCBP1’s precise role(s) in the HCV life cycle, we used assays to specifically examine different steps of the viral life cycle (**Figure 2**). Firstly, we explored whether PCBP1 knockdown had an effect on viral entry using the HCV pseudoparticle (HCVpp) system. HCVpp are lentiviral vectors with a firefly luciferase reporter gene pseudotyped with the HCV E1 and E2 glycoproteins (22). HCVpp engage with HCV-specific entry receptors, then enter cells by clathrin-mediated endocytosis; thus, to account for any changes in clathrin-mediated endocytosis, we used vesicular stomatitis virus (VSV) pseudoparticles (VSVpp) as a control. In addition, to verify that PCBP1 knockdown did not affect luciferase reporter gene expression, we assessed firefly luciferase expression from cells directly transfected with a FLuc reporter plasmid. In all cases, we found that depletion of endogenous PCBP1 had no impact on luciferase activity (**Figure 2A**). This suggests that PCBP1 knockdown does not affect FLuc reporter expression, clathrin-mediated endocytosis, or HCVpp entry.

**Figure 2.**
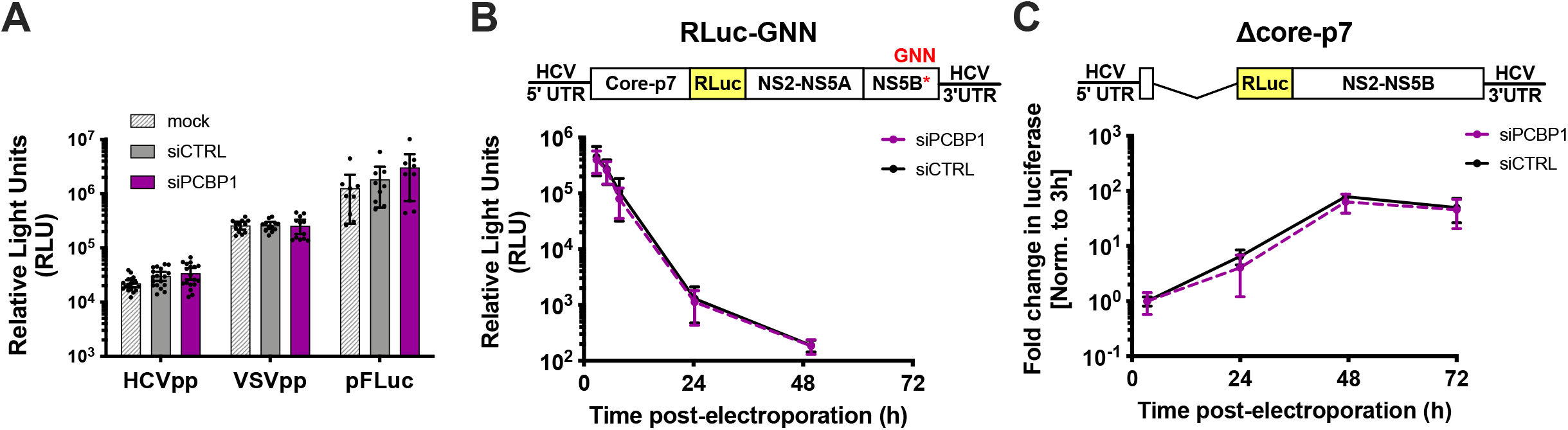
PCBP1 knockdown has no direct effect on HCV entry, translation, genome stability or RNA replication. **(A)** siRNA-transfected Huh-7.5 cells were spinoculated with luciferase reporter pseudoparticles expressing the HCV E1/E2 glycoproteins (HCVpp) or the VSV-G glycoprotein (VSVpp). In parallel, cells were transfected with a firefly luciferase expression plasmid. Samples were harvested 3 days post-infection/transfection and analyzed by luciferase assay. **(B)** siRNA-transfected Huh-7.5 cells were electroporated with replication-defective J6/JFH-1-RLuc-GNN RNAs, and luciferase activity was monitored at several timepoints in the two days post-electroporation. **(C)** Huh-7.5 cells were electroporated with the replication-competent Δcore-p7 J6/JFH RLuc RNA, and luciferase activity was monitored at several timepoints post-electroporation. RLuc values were normalized to the early timepoint (3h), to control for disparities in electroporation efficiency between experiments. All data are representative of at least three independent biological replicates and error bars represent the standard deviation of the mean.

To examine PCBP1’s impact on viral translation, we used a full-length J6/JFH-1 RLuc reporter RNA containing an inactivating mutation in the NS5B polymerase gene (GNN). The RLuc activity thus served as a direct measure of HCV IRES-mediated translation, and over time, this signal also served as a proxy measure for viral RNA stability. We found that the PCBP1 knockdown and control conditions had similar RLuc activity at all timepoints (**Figure 2B**). Moreover, the luciferase signal half-lives were nearly identical, with 2.68 h (95% CI 1.64 - 4.64) and 2.71 h (95% CI 1.43 - 5.50) for the PCBP1 knockdown and control conditions, respectively. Thus, PCBP1 knockdown does not directly impact either HCV IRES-mediated translation or genome stability.

To assess the impact of PCBP1 knockdown on HCV RNA replication, we made use of a J6/JFH-1 subgenomic replicon RLuc reporter RNA, with a deletion of the structural proteins (Δcore-p7), which allowed us to assess viral RNA replication in the absence of virion assembly (**Figure 2C**). We observed no significant differences in subgenomic replicon RNA accumulation during PCBP1 knockdown compared with control conditions, suggesting that PCBP1 knockdown does not affect viral RNA replication. Interestingly, we also observed no significant differences in luciferase activity during PCBP1 knockdown and control conditions of a full-length WT J6/JFH-1 RLuc genomic RNA or of a full-length J6/JFH-1 genomic RNA containing a GFP insertion in the NS5A gene (NS5A-GFP), which was previously shown to impair virion assembly without impairing viral RNA replication (**Supplementary Figure S3**) (6, 27). While this is seemingly in contrast to our findings with HCVcc, where PCBP1 depletion decreased viral RNA levels (**Figure 1C and D**); it is important to note that all of these reporter viral RNAs are defective in viral packaging due to the large RLuc reporter gene insertion, which results in >100-to >1000-fold reductions in infectious virion production (**Supplementary Figure S3**). Thus, taken together, our data suggests that PCBP1 knockdown does not have a direct effect on viral RNA replication.

### PCBP1 knockdown limits virus secretion

As we had found that silencing PCBP1 did not have a significant impact on HCV entry, translation, genome stability or viral RNA replication, we reasoned that PCBP1 was likely to be exerting an effect on virion assembly and/or secretion. To investigate this possibility, we used a nucleoside analog, 2’C-methyladenosine (2’CMA), to block viral RNA synthesis by the HCV NS5B RNA-dependent RNA polymerase and monitored virion production over time (**Figure 3**) (23). Similar to related studies using Zika virus, we reasoned that blocking viral RNA synthesis could allow us to assess if PCBP1 knockdown modulated viral assembly and secretion in the absence of genomic RNA production (28). To this end, we infected cells with JFH-1_T_ and, three days post-infection, treated cells with 2’CMA or vehicle (DMSO), and monitored viral RNA accumulation as well as intracellular and extracellular virion production over the next 12 hours (**Figure 3A**). In agreement with our previous findings, we observed an overall reduction in JFH-1_T_ viral RNA accumulation by day 3 post-infection during PCBP1 knockdown (0 h timepoint, **Figure 3B**). In addition, 2’CMA treatment efficiently blocked further viral RNA accumulation, which continued to increase under the vehicle (DMSO) condition (6-12 h timepoints, **Figure 3B**). We observed that intracellular viral titers were initially similar in both PCBP1 knockdown and control conditions, but that infectious virions accumulated at a slightly faster rate during PCBP1 knockdown in DMSO-treated cells (**Figure 3C and D**). In contrast, the 2’CMA treatment induced a similar decrease in intracellular titers during both PCBP1 knockdown and control conditions, resulting in similar rates of intracellular virion accumulation (**Figure 3C and D**). Interestingly, as we had observed previously (**Figure 1F**), PCBP1 knockdown resulted in an increase in extracellular titers which continued upon 2’CMA treatment (**Figure 3E and F**). Despite an overall reduction in extracellular virion production during 2’CMA treatment, we observed that PCBP1 knockdown still resulted in higher extracellular titers and virion secretion rates than the control condition under both 2’CMA and vehicle (DMSO) treatment (**Figure 3E and F**). Taken together, these results suggest that endogenous PCBP1 typically limits the secretion of infectious viral particles during the HCV life cycle.

**Figure 3.**
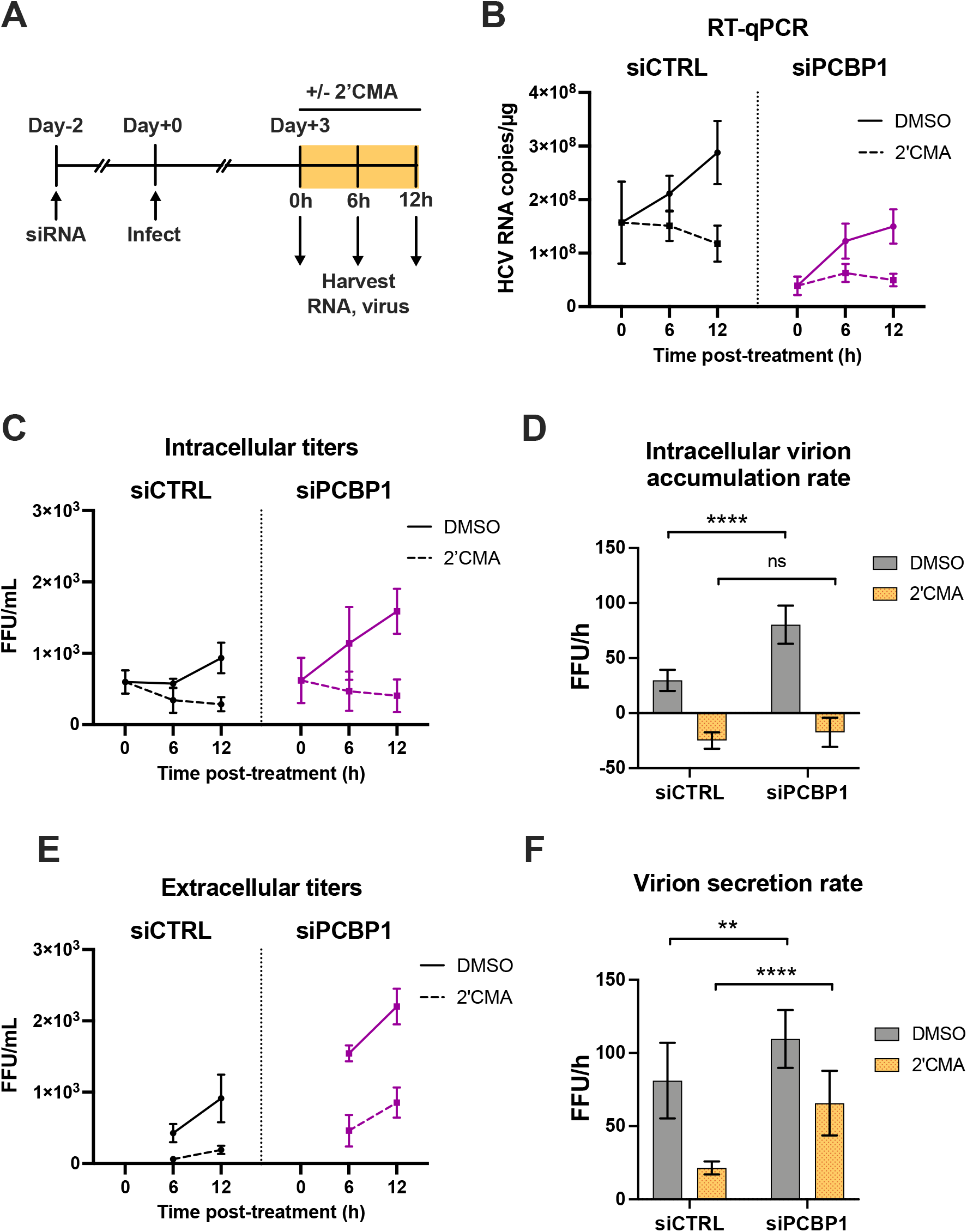
PCBP1 knockdown enhances infectious virion secretion even in the absence of RNA replication. **(A)** Schematic representation of the experimental approach for 2’CMA experiments: two days post-siRNA transfection, Huh-7.5 cells were infected with JFH-1_T_ (MOI = 0.05). Three days post-infection, the cell culture medium was replaced with media containing 2**’**CMA or DMSO (vehicle control). Total RNA, intracellular and extracellular virus were collected at t = 0, 6 and 12 h post-treatment. **(B)** Quantitative RT-PCR analysis after 2’CMA treatment. **(C)** Intracellular viral titers and **(D**) intracellular virus accumulation rate calculated by linear regression. **(E)** Extracellular viral titers and **(F)** virus secretion rates calculated by linear regression. All data are representative of three independent replicates and error bars in (B), (C) and (E) represent the standard deviation of the mean. Error bars in (D) and (F) represent the slopes of the linear regressions ± standard error. Statistical significance was calculated by two-way ANOVA. (ns, not significant; ** p < 0.01; **** p < 0.0001).

### Blocking virion secretion partially restores HCV RNA accumulation during PCBP1 knockdown

During virion secretion, viral genomes that have been packaged into new infectious particles are removed from the cell. Since our previous findings suggested that PCBP1 knockdown increases virion secretion, we wondered if this increased egress of assembled viral particles (packaged viral genomic RNAs) might explain our initial observation that PCBP1 knockdown led to an overall reduction in viral RNA accumulation (**Figure 1C-D**). To directly test this, we treated HCV-infected cells with Brefeldin A (BFA), a protein trafficking inhibitor previously shown to prevent HCV virion secretion (**Figure 4**) (24). As previously reported, we found that BFA treatment blocked the secretion of infectious virions, resulting in a sharp decline in extracellular titers and an increase in intracellular titers relative to untreated cells or to vehicle (DMSO) treatment (**Figure 4A and B**). Consistent with our prior results, PCBP1 knockdown resulted in a greater than 2-fold decrease in viral RNA accumulation, while BFA treatment resulted in a partial rescue of HCV RNA accumulation during PCBP1 knockdown (**Figure 4C**). Thus, blocking cellular secretory pathways, thereby preventing virion secretion, can partially rescue intracellular viral RNA accumulation defects resultant from PCBP1 knockdown.

**Figure 4.**
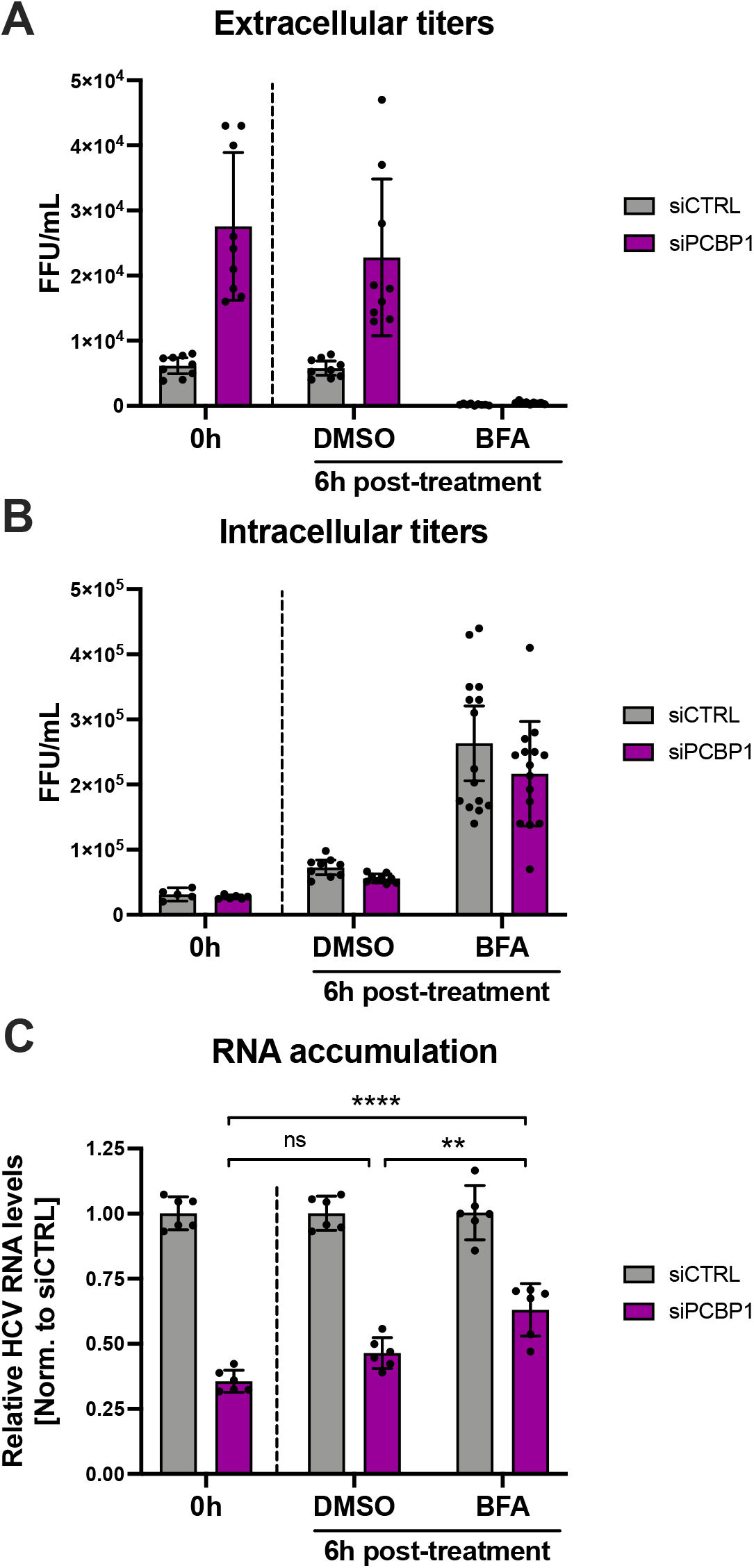
Blocking HCV secretion partially rescues intracellular RNA accumulation during PCBP1 knockdown. Three days post-JFH-1_T_ infection (MOI 0.05), the cell culture medium was replaced with media containing Brefeldin A (BFA) or DMSO (vehicle control). Total intracellular RNA, intracellular virus, and extracellular virus were collected at 0 h and 6 h post-treatment. **(A)** Extracellular (secreted) viral titers and **(B)** intracellular viral titers from untreated cells (0h) and after 6h of DMSO or BFA treatment, quantified by FFU assay. **(C)** Intracellular RNA accumulation relative to the siCTRL condition, assessed by RT-qPCR analysis. All data are representative of three independent biological replicates, and error bars represent the standard deviation of the mean. Statistical significance was calculated by paired t-test (ns, not significant; ** p < 0.01; **** p < 0.0001).

## DISCUSSION

Herein, we investigated the role of PCBP1 in the HCV life cycle. Previous studies had found that PCBP1 interacts with the HCV 5’ UTR *in vitro* (10, 15), and that PCBP1 silencing inhibited HCV RNA accumulation in cell culture (16), yet the precise steps of the HCV life cycle modulated by PCBP1 remained unclear. Our systematic evaluation of the effects of PCBP1 knockdown on individual steps of the viral life cycle revealed that PCBP1 depletion did not directly affect virus entry, translation, genome stability or viral RNA replication, but rather resulted in increased infectious virion production in cell culture. Specifically, PCBP1 knockdown significantly increased virion secretion, resulting in the depletion of intracellular viral RNA. Additionally, inhibiting virion secretion could partially restore HCV RNA accumulation during PCBP1 knockdown. Thus, our findings suggest a model whereby endogenous PCBP1 normally limits virion assembly and secretion, such that viral genomes remain available to engage in the translation and replication steps of the viral life cycle (**Figure 5**). As such, PCBP1 depletion increases the proportion of viral genomes engaged in virion assembly and secretion, which decreases the pool of translating/replicating viral RNAs. We anticipate that these differences in intracellular viral RNA levels would compound over time, with PCBP1-replete cells rapidly increasing their pool of intracellular viral RNA through repeated cycles of translation and RNA replication, while the smaller pool of translating/replicating RNAs in PCBP1-depleted cells would result in a much slower rate of viral RNA accumulation over time.

**Figure 5.**
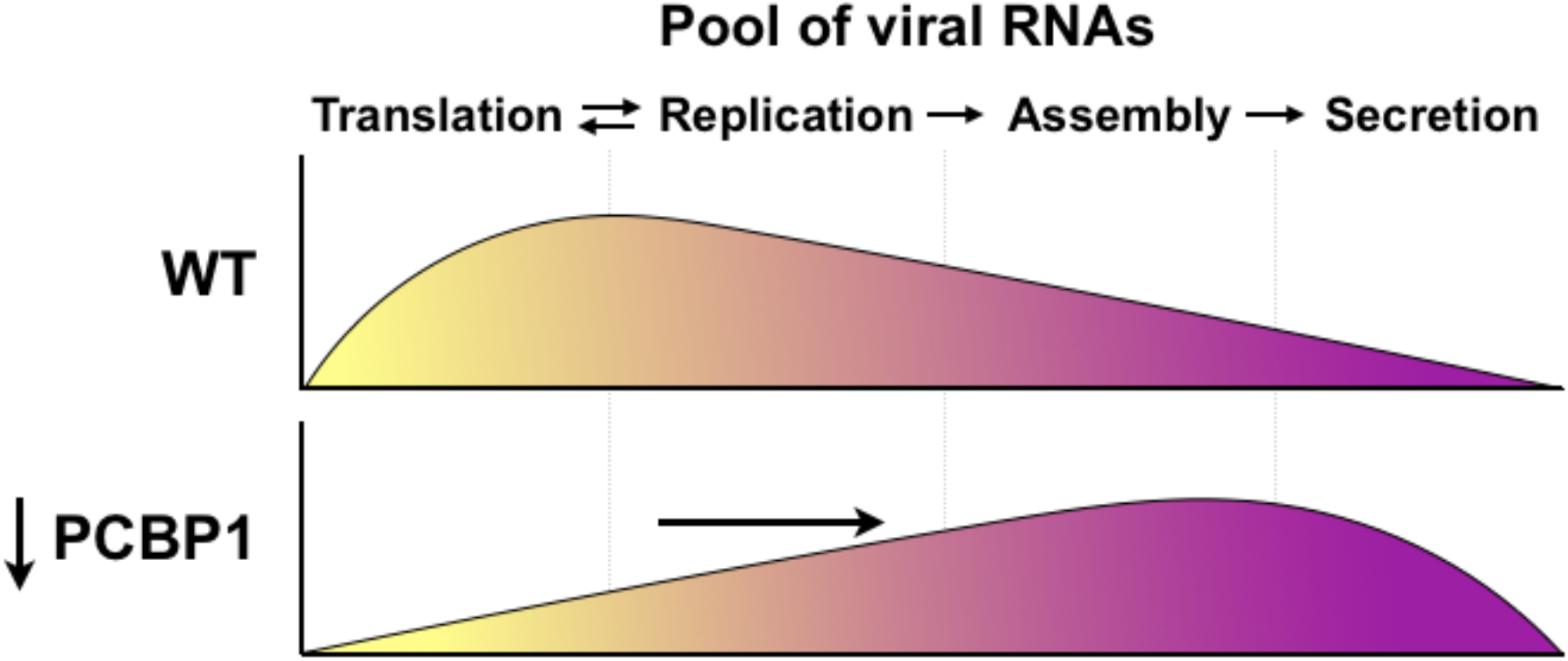
Model for PCBP1’s effects on HCV assembly and secretion. Under wild-type conditions, endogenous PCBP1 limits HCV assembly and secretion, and the majority of intracellular viral RNAs are engaged in translation and viral genome replication. In contrast, PCBP1 knockdown promotes virion assembly and secretion. This results in a shift in viral RNA utilization such that an increased proportion of viral RNAs are engaged in viral particle production, leaving a smaller proportion of viral RNAs to engage in the translating/replicating pool.

Notably, most of the experiments conducted herein made use of JFH-1_T_, a cell culture-adapted strain that produces greater yields of infectious virions in cell culture than the parental JFH-1 strain (5, 17). To verify that the phenotype we observed was not an artefact of the JFH-1_T_ strain itself, we validated our findings with J6/JFH-1, a chimeric virus that is widely used in HCVcc studies, commonly referred to as Jc1 (18). Knockdown of PCBP1 had a similar effect on JFH-1_T_ and J6/JFH-1 protein expression, RNA accumulation, and extracellular viral titers; however, we found that while JFH-1_T_ intracellular titers were not significantly different from control condition, J6/JFH-1 intracellular titers were significantly decreased by approximately 2.6-fold (**Figure 1E**). Nonetheless, PCBP1 knockdown still resulted in an overall greater than 2-fold increase in the total number of infectious virions produced by both strains (**Supplementary Figure S1**). Since an infectious viral particle must have undergone the process of virion assembly, this suggests that PCBP1 depletion enhances the assembly of infectious virions in addition to their subsequent secretion. Importantly, this data could reflect pre-existing strain differences in virion assembly, as J6/JFH-1 has previously been reported to be more efficient than JFH-1 in virion assembly (29, 30). Specifically, the J6 core protein was shown to localize closer to the sites of virion assembly on the endoplasmic reticulum (ER) than the JFH-1 core protein, which accumulates to a greater extent on lipid droplets (29, 30). Furthermore, J6/JFH-1 allocates a greater proportion of its viral genomes to virion assembly than JFH-1, which allocates more viral RNAs to the translating/replicating pool (29, 30). Thus, it is possible that PCBP1 knockdown promotes both the assembly and secretion of J6/JFH-1 virions, but minor gains in the already efficient virion assembly step are less apparent than the gains in virion secretion for this particular strain.

Interestingly, PCBP1 has also been previously implicated in attenuation of antiviral signalling, specifically in the regulation and turnover of MAVS, a signal transduction protein directly downstream of RIG-I, which is known to limit HCV infection (31). However, all of our conclusions were drawn from experiments conducted in the Huh-7.5 cell line, which has well-documented defects in the RIG-I antiviral signalling pathway (32). Additionally, our own brief explorations of interferon induction and MAVS turnover did not reveal any significant differences between PCBP1 knockdown and control conditions (data not shown). Moreover, the HCV NS3-4A protease also inactivates MAVS during infection to reduce antiviral signaling and recognition of the viral RNA (33, 34). However, as hepatocytes typically express RIG-I and MAVS, we cannot rule out the possibility that PCBP1 may also modulate this pathway during infection *in vivo*.

Finally, PCBP1’s closely related paralogs, hnRNP K and PCBP2, have been more extensively studied during HCV infection. While the hnRNP K protein was shown to restrict infectious virion production, the PCBP2 protein has been suggested to modulate viral translation and RNA replication (13, 14). Since the PCBP1 amino acid identity is far more similar to PCBP2 (~80% identity) than hnRNP K (~24% identity), we were initially surprised that our findings for PCBP1 in HCV infection closely matched those reported for hnRNP K. Yet, while PCBP1 and PCBP2 have been shown to perform similar functions, they have also been reported to play distinct roles during poliovirus, vesicular stomatitis virus, and human immunodeficiency virus infection (35–37). In addition to the similarities we observed between PCBP1 and the effects previously reported for hnRNP K, our results and conclusions also echo those reported for the IGF2BP2/YBX-1 complex, METTL3/METTL14 *N*6-methyladenosine (m^6^A) writers, and the YTHDF m^6^A-binding proteins during HCV infection (13, 38, 39). All of these complexes have been reported to inhibit HCV infectious particle production, with no effect on viral translation or RNA replication - with the exception of the IGF2BP2/YBX-1 complex, which plays an additional role in facilitating viral RNA replication (38). It is currently unclear if PCBP1, hnRNP K, the YBX-1 complex, or m^6^A modification of the HCV genome inhibit virion assembly as components of a common pathway, or through distinct and/or additive mechanisms. Interestingly, high-throughput affinity capture and proximity ligation studies have found that PCBP1 interacts with hnRNP K, IGF2BP2, YBX-1, METTL3 and METTL14; although this has been demonstrated only in non-hepatic cell lines to date (40–42). Should these interactions be conserved during HCV infection, it seems plausible that PCBP1 may participate in one or more of these complexes which together inhibit HCV virion assembly and secretion. Future investigations will thus be needed to reveal whether these proteins function in an overlapping or distinct manner; and, are likely to further improve our understanding of HCV virion assembly as well as how this process is regulated by cellular RNA binding proteins.

In conclusion, our results support a model whereby endogenous PCBP1 limits HCV infectious particle production. By preventing virion assembly and secretion, PCBP1 indirectly enhances viral RNA accumulation. The model presented herein helps to inform our understanding of how cellular RNA-binding proteins modulate HCV genomic RNA utilization during the viral life cycle, specifically as it pertains to virion assembly. While the precise molecular mechanism(s) employed by PCBP1 to inhibit HCV assembly and egress remain to be characterized, similar phenotypes reported for hnRNP K, IGF2BP2/YBX-1 and m^6^A modification of the HCV genome offer promising leads for future investigations.

## ACKNOWLEDGEMENTS

We would like to acknowledge Charlie Rice (Rockefeller University) for kindly providing the Huh-7.5 cells, pJ6/JFH (pJc1), pJ6/JFH FL RLuc WT and GNN plasmids; Rodney Russell (Memorial University) for providing JFH-1_T_; Mamata Panigrahi and Joyce Wilson (University of Saskatchewan) for the pJ6/JFH mono RLuc NS2 (“Δcore-p7”) plasmid; Martin J Richer (McGill University) for the 293T cells; John Law (University of Alberta) for the kind gift of HCVpp. We are also grateful to Nathan Taylor, Julie Magnus, and Carolina Camargo (McGill University) for technical support, and to Craig McCormick’s graduate journal club (Dalhousie University) for their critical feedback on a previous iteration of this manuscript.

## AUTHOR CONTRIBUTIONS

S.E.C. and S.M.S. designed the study; S.E.C. and M.R. performed the experiments and analyzed the data, and S.E.C and S.M.S. wrote and edited the manuscript.

## SUPPLEMENTARY INFORMATION

### SUPPLEMENTARY METHODS

#### Plasmids

To generate the pPCBP1(N)FLAG-puro plasmid, the complete PCBP1 coding sequence was amplified using PCBP1noAUG-KpnI-FW (5’-AAG CTG GCG GTA CCG GAG ATG CCG GTG TGA CTG-3’) and PCBP1-XhoI-RV (5’-TTT TTC TC GAG CTA GCT GCA CCC CAT GCC CTT-3’) and the high-fidelity Hot Start Q5 DNA polymerase (NEB). The PCR product was digested with KpnI and XhoI and ligated in a pcDNA3.1(+)-derived plasmid that contained a 3xFLAG tag located downstream of a human cytomegalovirus (CMV) immediate early enhancer/promoter and upstream of a bovine growth hormone (bGH) polyadenylation signal; this insertion left the PCBP1 coding sequence in frame with the upstream 3x FLAG tag. To note, this pcDNA3.1(+)-derived vector also contained a puromycin resistance gene located downstream of a phosphoglycerate kinase 1 (PGK) promoter, and upstream of a simian virus 40 (SV40) polyadenylation signal.

To generate the pFLuc-puro plasmid, the complete Firefly luciferase coding sequence was amplified using FLuc-NheI-FW (5’-ATC CGC TAG CAT GGA TTA CAA GGA C-3’) and FLuc-XhoI-RV (5’-AAG GTA TCT CGA GTT AGT AAA CAA GAT AAT TGC TCC TAA AGT A-3’) using the high-fidelity Hot Start Q5 DNA polymerase (NEB). The PCR product was digested with NheI and XhoI and ligated into the same pcDNA3.1(+)-derived plasmid, between the CMV enhancer/promoter and the bGH polyadenylation signal; this insertion removed the 3xFLAG tag.

The pJ6/JFH1 FL RLuc WT (“RLuc-wt”) viral sequence contains include a *Renilla* luciferase (RLuc) reporter gene inserted between p7 and NS2 (1). The pJ6/JFH-1 FL RLuc-NS5A-GFP (“NS5A-GFP”) plasmid contains a GFP insertion between P2390 and L2391 within the NS5A domain III, as previously described (2). The NS5A-GFP region was subcloned into the pJ6/JFH-1 FL RLuc WT backbone using the *AvrII* and *XbaI* restriction sites.

To construct a JFH-1_T_-RLuc viral genome template, a MluI site was created between the p7 and NS2 genes by PCR amplification of the pJFH-1_T_ plasmid using JFH1-p7NS2-MluI-FW (5’-CCT ATG ACG CGT CTG TGC ACG GAC AGA TAG GC-3’), JFH1-p7NS2-MluI-RV (5’-CAG ACG CGT CAT AGG CAT AAG CCT GCC G-3’), and the high-fidelity Hot Start Q5 DNA polymerase (NEB). The PCR product was digested with MluI and dephosphorylated with calf intestinal phosphatase (Quick CIP, NEB) and ligated with an insert containing a *Renilla* luciferase (RLuc) with a C-terminal foot-and-mouth disease virus 2A peptide (FMDV2A), which had been cut out of the pJ6/JFH WT RLuc plasmid by MluI digestion. The correct orientation of the insert was screened by digestion with BstBI and NotI and by sequencing. Once a plasmid with the correct insert orientation was identified, the E1 through NS2 region was cut out using the BsiWI and NotI unique cut sites and ligated into the original pJFH-1_T_ plasmid backbone. The inserted E1 through NS2 region was sequenced to verify that the PCR amplifications had not introduced any unexpected substitutions.

#### Puromycin selection of cells and infection with JFH-1_T_

To select cells that would stably express PCBP1(N)FLAG or the FLuc control protein, 15-cm dishes were seeded with 2 x 10^6^ Huh-7.5 cells one day prior to transfection with 36 μg of pPCBP1(N)FLAG-puro or pFLuc-puro plasmid and 30 μL Lipofectamine 2000 diluted in 12 mL Opti-MEM serum-free media (Thermo Fisher Scientific). Two days post-transfection, the media on all transfected plates and one non-transfected plate was changed for complete Huh-7.5 media supplemented with 3 μg/mL puromycin (Thermo Fisher Scientific), which was replenished every two to three days, passaging the cells as necessary. Once all cells in the non-transfected plate were completely dead (within 7 days of selection), selected cells could be used for infection experiments. Selected cells were transfected with siRNA duplexes and infected with JFH-1_T_ at an MOI of 0.05 as described in the article’s main methods; the selection pressure of 3 μg/mL puromycin was maintained in these cells’ media until their infection with JFH-1_T_, after which they were kept in puromycin-free media.

## SUPPLEMENTARY FIGURES

**Supplementary Figure S1.**
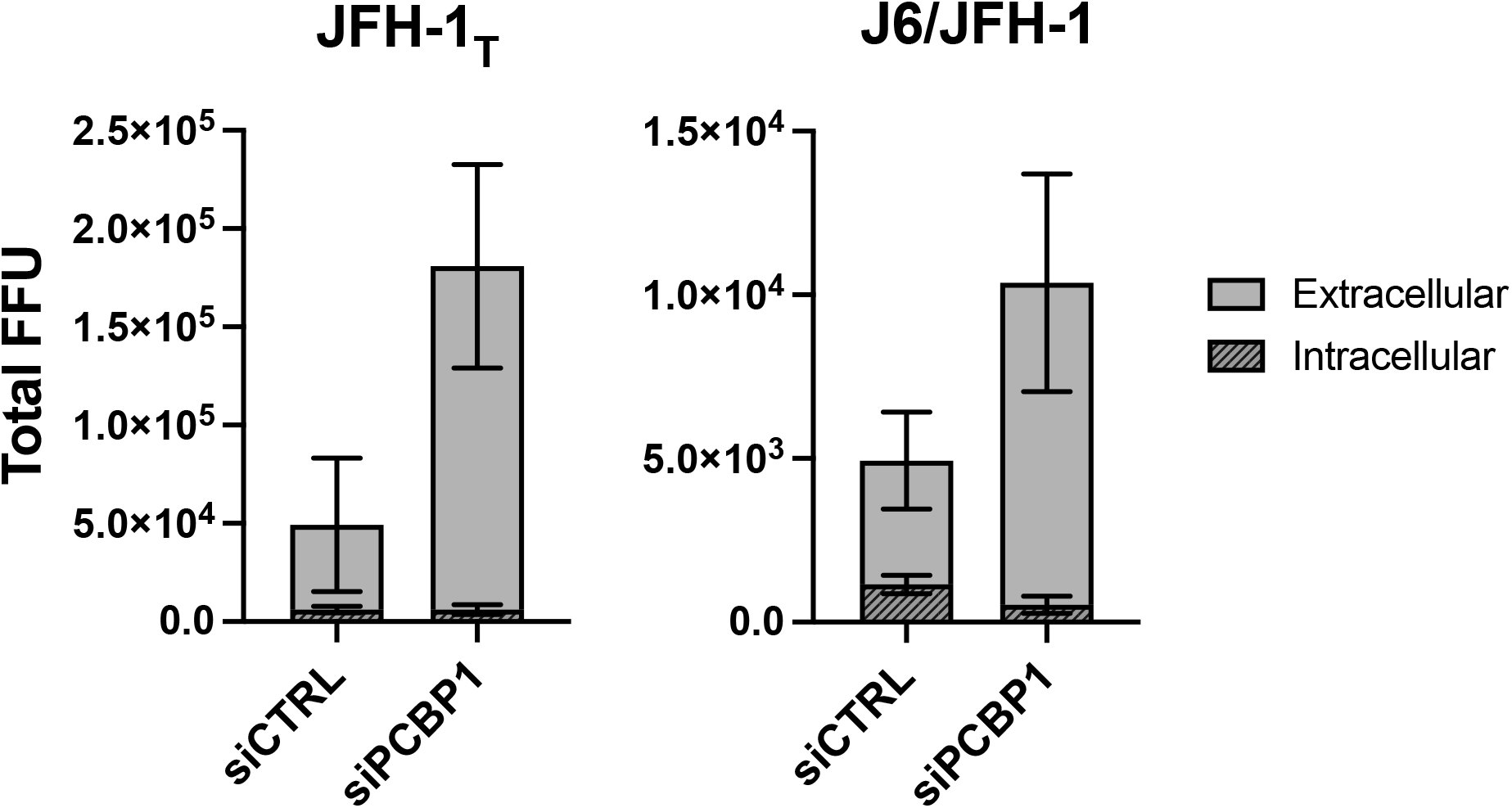
PCBP1 knockdown increases the total quantity of infectious virions produced by the JFH-1_T_ and J6/JFH-1 strains. To calculate the total quantity of infectious virions (FFU) present in a sample, the viral titer (FFU/mL) was multiplied by the total volume of the sample (mL). These data were derived from the same experiments presented in Figure 1 and are representative of three independent biological replicates. Error bars represent the standard deviation of the mean.

**Supplementary Figure S2.**
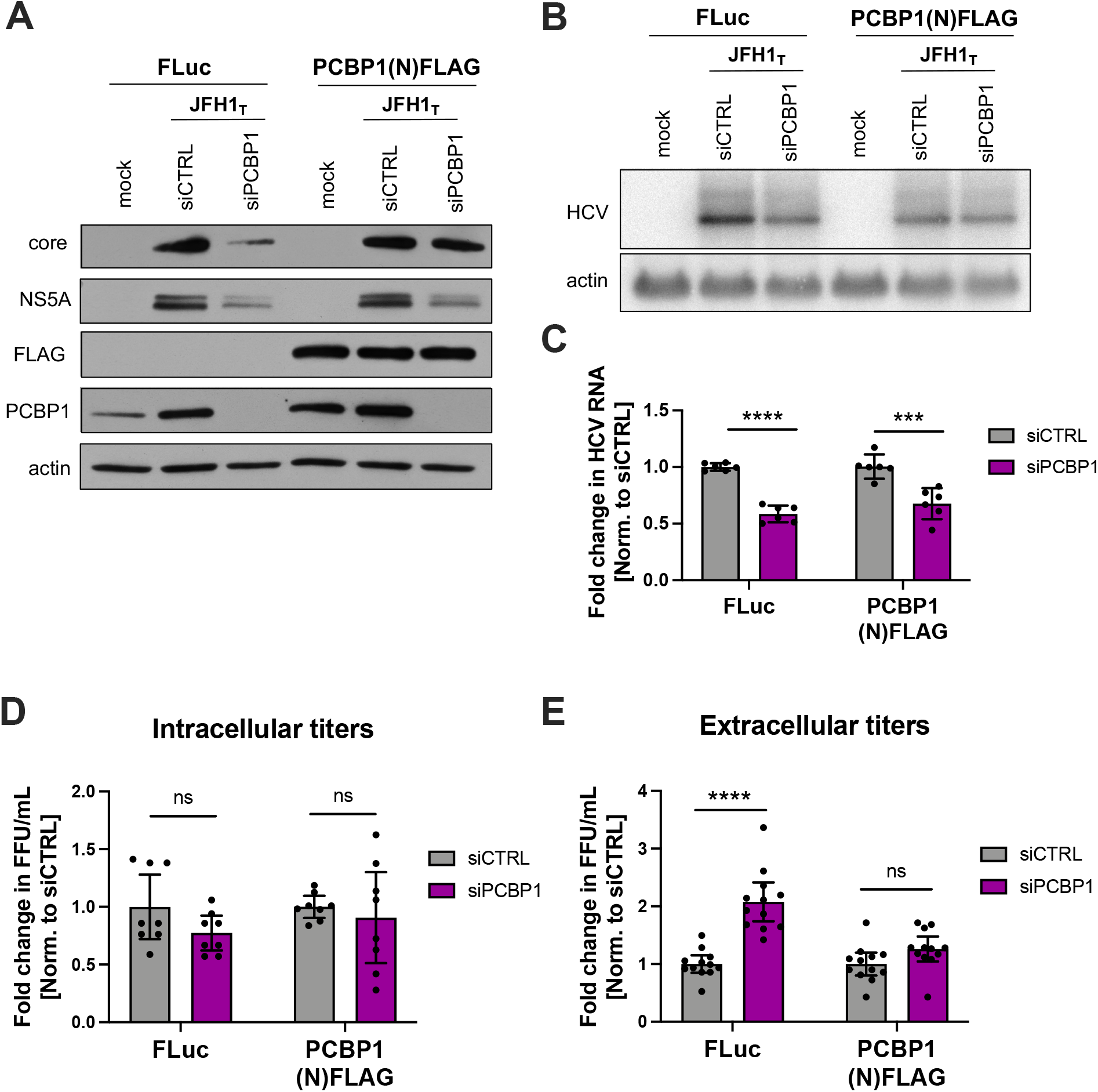
Ectopic PCBP1 expression can reduce the impact of endogenous PCBP1 knockdown on extracellular viral titers. Huh-7.5 cells that were puromycin-selected to stably express ectopic Firefly luciferase (FLuc, control) or PCBP1 with a N-terminal FLAG tag (PCBP1(N)FLAG) were transfected with siCTRL or siPCBP1 two days prior to infection with JFH-1_T_ (MOI 0.05). Three days post-infection, total intracellular protein, RNA, and intracellular and extracellular virus were collected. **(A)** Viral protein expression analysis by Western blot. **(B)** Viral RNA accumulation analysis by Northern blot and **(C)** quantification by RT-qPCR. **(D)** Intracellular and **(E)** extracellular (secreted) virus titers, quantified by FFU assay. For (C-E), data from each experiment was normalized to the mean of the cell-matched siCTRL condition before replicate data were combined. All data are representative of three independent biological replicates, with the exception of (B) which is an N = 1. Error bars represent the standard deviation of the mean. Statistical significance was calculated by paired t-test (ns, not significant; *** p < 0.001; **** p < 0.0001).

**Supplementary Figure S3.**
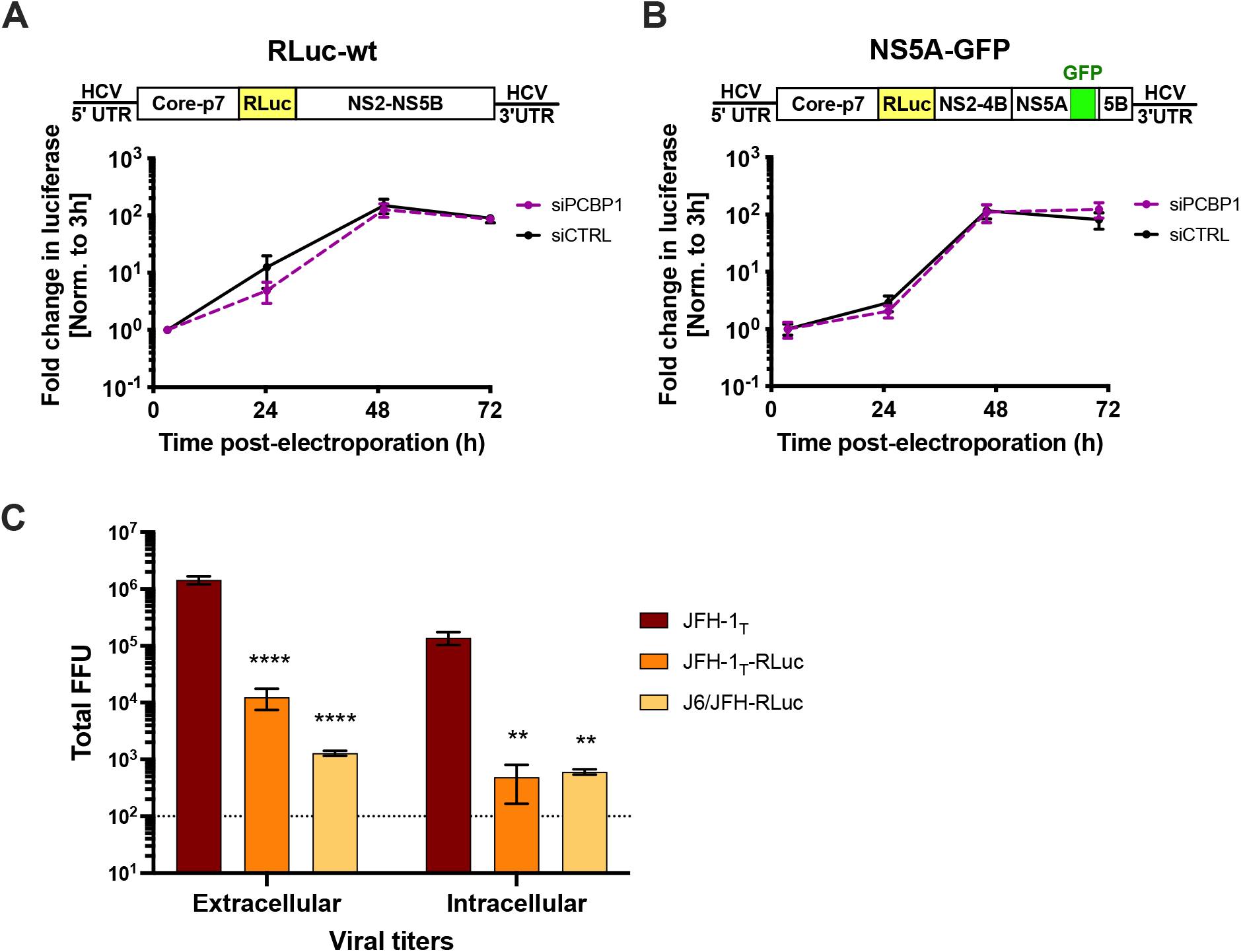
PCBP1 has no effect on RNA replication in the absence of efficient virion assembly, and the addition of a RLuc gene reduces the assembly efficiency of HCV genomes. SiRNA-transfected Huh-7.5 cells were electroporated with 5 μg of (**A**) full-length J6/JFH RLuc WT RNA, or of (**B**) a full-length J6/JFH RLuc with a GFP insertion in the NS5A gene (NS5A-GFP), which had previously been shown to impair virion assembly without impairing viral RNA replication. Luciferase activity was monitored for three days post-electroporation, and RLuc values were normalized to the early timepoint (3h) to control for disparities in electroporation efficiency between experiments. (**C**) Equal amounts (10 μg) of JFH-1_T_, JFH-1_T_-RLuc, or J6/JFH-RLuc RNAs were electroporated into Huh-7.5 cells; intracellular and extracellular viruses were collected three days post-electroporation and titered by focus-forming unit assay. Compared with the untagged JFH-1_T_ RNA electroporations, the extracellular titers were reduced by over 100-fold and 1000-fold for JFH1_T_-RLuc and J6/JFH-RLuc, respectively; intracellular titers were reduced by more than 200-fold for both luciferase reporter genomes. Data in (A) and (B) are representative of three independent biological replicates; data in (C) is representative of two independent biological replicates, and the limit of detection is indicated. Error bars represent the standard deviation of the mean. Statistical significance was calculated by t-test (** p < 0.005, **** p < 0.0001)

